# Inter-species microbiota transplantation recapitulates microbial acquisition and persistence in mosquitoes

**DOI:** 10.1101/2021.10.06.463328

**Authors:** Kerri L. Coon, Shivanand Hegde, Grant L. Hughes

## Abstract

**Background:** Mosquitoes harbor microbial communities that play important roles in their growth, survival, reproduction, and ability to transmit human pathogens. Microbiome transplantation approaches are often used to study host-microbe interactions and identify microbial taxa and assemblages associated with health or disease. However, no such approaches have been developed to manipulate the microbiota of mosquitoes.

**Results:** Here, we developed an approach to transfer entire microbial communities between mosquito cohorts. We undertook transfers between (*Culex quinquefasciatus* to *Aedes aegypti)* and within (*Ae. aegypti to Ae. aegypti*) species to validate the approach and determine the number of mosquitoes required to prepare donor microbiota. After the transfer, we monitored mosquito development and microbiota dynamics throughout the life cycle. Typical holometabolous lifestyle-related microbiota structures were observed, with higher dynamics of microbial structures in larval stages, including the larval water, and less diversity in adults. Microbiota diversity in recipient adults was also more similar to the microbiota diversity in donor adults.

**Conclusions:** This study provides the first evidence for successful microbiome transplantation in mosquitoes. Our results highlight the value of such methods for studying mosquito-microbe interactions and lay the foundation for future studies to elucidate the factors underlying microbiota acquisition, assembly, and function in mosquitoes under controlled conditions.

## Background

A substantial body of evidence has emerged revealing the importance of microbiota to the biology of the animal hosts they associate with, which has stimulated broad interest in understanding the assembly of these communities. However, the daunting complexity of the microbiota present in many higher eukaryotes and the lack of conventional microbiology techniques to culture these microbes has limited our ability to address important questions in the field. As such, the mechanisms facilitating host-microbe interactions and the functional role of the microbiome as a holistic unit are poorly elucidated. Microbiota transplantation approaches are one promising technique for basic research and therapeutics, but studies employing these techniques have mainly been undertaken in mammalian systems [1-6] and there has been little attempt to transfer these approaches to medically or agriculturally important insects.

Microbiome research has expanded in mosquitoes given their medical relevance and interesting biology. As holometabolous insects, mosquitoes have distinct aquatic and terrestrial life stages, including larvae that molt through four aquatic instars before pupating on the water’s surface and emerging as terrestrial adults [7, 8]. All of these stages harbor gut microbial communities dominated by bacteria that can vary tremendously in diversity and abundance over time and space [9]. Larvae acquire their gut microbiota from their aquatic environment during feeding [10-14], after which bacteria experience periods of extreme turnover as parts of the larval gut are continuously shed and replaced during feeding, molting, and metamorphosis to the adult stage [15, 16]. The adult gut, in contrast, is initially seeded by bacteria from larvae and/or the larval environment but thereafter may be modulated by adult sugar and blood feeding activities [9], the latter of which underlies the ability of adult female mosquitoes to acquire and transmit disease-causing pathogens to humans [17]. While most of the larval and adult mosquito microbiota is thought to be restricted to the midgut [18-21], bacteria and other microbes are also known to colonize other mosquito tissues [22-25]. These include the common bacterial endosymbiont *Wolbachia*, which infects the germline of *Culex quinquefasciatus*, but not *Aedes aegypti*, and is transovarially transmitted to offspring each generation [26].

Altogether, the microbiota associated with the mosquito gut and other tissues can have profound impacts on mosquito biology by modulating larval growth and development [10, 12, 27-29] as well as adult survival [30-32], reproduction [27, 33], and the competency of adult female mosquitoes to transmit human pathogens [34]. As such, there is a growing interest in exploiting microbes for vector control [35-37]. However, available tools to manipulate the microbiota in mosquitoes are limited, lagging behind research in other systems [38-41]. Antibiotic treatment has been commonly used to perturb microbiota for experiments investigating tripartite interactions between mosquitoes, their microbiota, and human pathogens. While these experiments have provided insights into the role of bacteria in mosquito vector competence [34, 42-44], the use of antibiotics has its limitations. For example, some bacteria may not be susceptible to the antibiotics, meaning these approaches alter the microbiota rather than eliminate all microbiota [12, 45]. Furthermore, antibiotics can have off-target effects on host physiology and can affect mitochondria [46-49]. Alternatively, introduction of specific bacterial taxa into mosquito larvae or adults can be achieved by spiking the larval water or sugar solution, respectively [50-52]. However, while these approaches are effective at transferring specific bacterial taxa to mosquitoes, they likely do not reflect acquisition processes that occur in the field and can only be undertaken with culturable microbes.

We previously developed an approach to generate and maintain gnotobiotic mosquitoes colonized by individual bacterial taxa by sterilizing the surface of mosquito eggs and inoculating water containing bacteria-free (axenic) larvae hatched from surface-sterilized eggs under sterile conditions [10]. Similarly, the generation of axenic adult mosquitoes that can thereafter be inoculated with individual bacterial taxa via a sugar meal has been undertaken using heat-killed bacteria, supplementation of axenic larval cultures with eukaryotes, or reversible colonization [28, 53-55]. While these approaches have been used to broadly examine the biology of mosquito-microbe interactions and decipher the role of bacterial microbiota on host biology [10, 12, 21, 27, 28, 56-58], expanding these techniques to transfer complete or tailored microbial communities would be highly desirable.

Here, we developed an approach to transfer entire microbial communities between mosquito cohorts. We undertook transfers between (*Cx. quinquefasciatus* to *Ae. aegypti)* and within (*Ae. aegypti to Ae. aegypti*) species to validate the approach and determine the number of mosquitoes required to prepare donor microbiota. After the transfer, we monitored mosquito development and microbiota dynamics throughout the life cycle. Typical holometabolous lifestyle-related microbiota structures were observed, with higher dynamics of microbial structures in larval stages, including the larval water, and less diversity in adults. Furthermore, the diversity of microbiota present in recipient adults was more similar to the microbiota present in donor adults. Altogether, our results effectively demonstrate the transfer of a complete microbiota from one mosquito cohort to another and lay the foundation for future studies to examine microbiota assembly and function in mosquitoes colonized by defined microbiomes with potential for exploitation for mosquito and disease control.

## Methods

### Donor sample collection and preparation of donor pools for microbiota transplantation

*Ae. aegypti* (Galveston) and *Cx. quinquefasciatus* (Houston) mosquitoes used as donors in this study were reared under conventional conditions as described previously [57]. In order to characterize microbiota diversity in both donor species, 150 individual three-to-four-day-old sugar-fed adult females of each species were collected, surface sterilized by immersing in 70% ethanol for five minutes followed by three five minute washes in sterile 1X PBS, and stored at −20°C for downstream sequencing. Adult mosquitoes from both species were also used to generate donor pools for microbiota transplantation. In brief, cohorts of 10, 20, 40, or 80 three-to-four-day-old adult female mosquitoes were collected, surface-sterilized in 70% ethanol for five minutes, rinsed three times in sterile 1X PBS for five minutes, and transferred individually to sterile 2 ml safe lock tubes (Eppendorf, Hamburg, Germany) containing 5 mm steel beads and 500 µl of sterile 1X PBS (Fig. 1). Tubes were then homogenized at a frequency of 26 Hz/sec for one minute, briefly centrifuged to collect debris, and 50 µl of the homogenate from each tube was pooled (Fig. 1). Pooled homogenates were subsequently centrifuged at 5000 × g for five minutes to pellet any remaining debris, and the resulting supernatant was passed through a 5 µm filter to produce a final filtrate for transplantation. Filtrates, which ranged in volume from 500 µl (pool of 10) to 4 ml (pool of 80), were finally adjusted to a total volume of 50 ml using sterile water, and we repeated the entire process a total of three times to produce a total of 150 ml of each donor pool prior to use in downstream experiments (Fig. 1). A 200 µl aliquot of each pool was also retained and immediately subjected to genomic DNA isolation using a NucleoSpin Tissue Kit (Machery-Nagel, Düren, Germany). The resulting DNA was then stored at −20°C for downstream sequencing.

**Figure 1.**
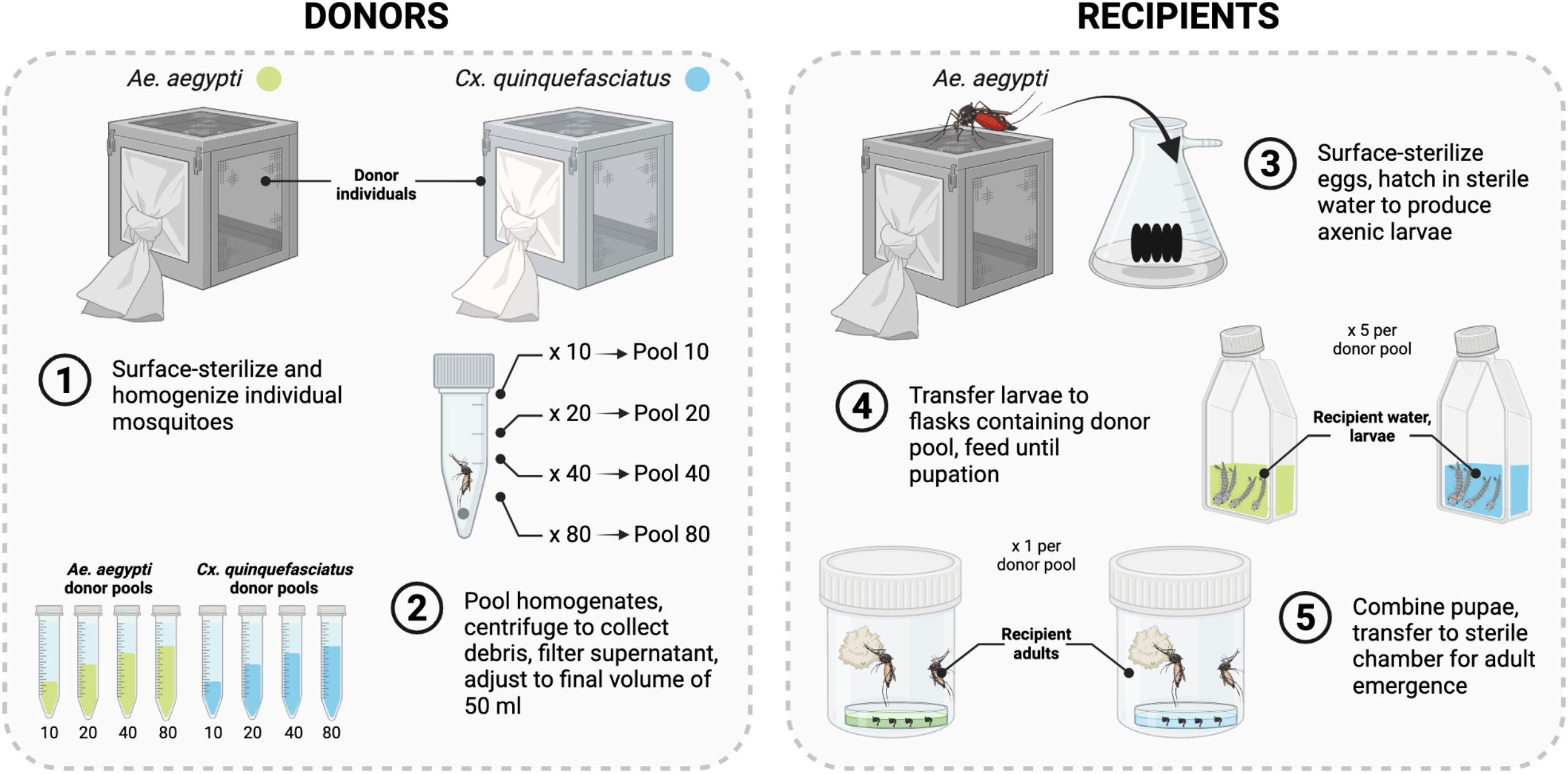
Overview of methodology used to perform inter- and intra-species microbiota transplantations in mosquitoes. (Left) Preparation of *Ae. aegypti* and *Cx. quinquefasciatus* donor pools. Individual 3-4-day-old sugar-fed adult females from our standard laboratory colonies were collected, surface-sterilized, and homogenized (1). Individual homogenates were then pooled and centrifuged to collect debris prior to filtering of the resulting supernatant to produce a final filtrate (2). Filtrates, which ranged in volume from 500 µl (Pool 10) to 4 ml (Pool 80), were then adjusted to a total volume of 50 ml using sterile water prior to transplantation (2). (Right) Transplantation of donor pools into a focal host species (*Ae. aegypti*). Eggs laid by blood-fed adult females from the standard laboratory colony were surface-sterilized and hatched in sterile water to produce axenic larvae (3). Larvae were then transferred to replicate flasks (*n*=5) containing a 50 ml suspension of a given donor pool and provided sterilized diet every other day until pupation (4). Pupae produced from replicate flasks containing the same donor pool were finally pooled in water from the larval rearing flasks and transferred to a sterile plastic chamber for adult emergence (5). Donor and recipient samples collected for sequencing are indicated in bold. See *Methods* for more information. Created with BioRender.com.

### Microbiota transplantation and recipient sample collection

Axenic (microbe-free) *Ae. aegypti* (Galveston) L1 larvae served as the recipient host for all transplantations. In brief, axenic larvae were prepared by submerging eggs in 70% ethanol for 5 minutes, then transferring to a solution of 3% bleach and 0.01% Decon-Quat 200V (Ab Scientific Ltd) for 3 minutes, followed by a wash in 70% ethanol for 5 minutes, and finally rinsing three times in sterile water (Fig. 1). Eggs were then transferred to sterile water and vacuum hatched in sterile containers. Twenty axenic first instar larvae were then transferred to each of five T75 vented tissue culture flasks (Thermo Fisher Scientific, Waltham, MA USA) containing 50 ml suspension of a given donor pool (described above) and 60 µl of autoclaved powdered TetraMin pellets (Tetra, Melle, Germany) reconstituted with sterile water to a final concentration of 1 mg/ml (Fig. 1). In addition to the donor treatments, a negative control was done whereby larvae were transferred to flasks containing sterile water and diet only (*i*.*e*., no microbes) and maintained alongside experimental flasks. All larvae (control and experimental) were provided sterilized diet every two days until the treatment groups pupated (control group did not pupate and died at L2 or L3) while water from control flasks was used to screen for contamination throughout the experiment as described previously [10]. Flasks were maintained at 70% humidity and 27°C.

In order to assess successful microbiome transplantation, four sets of recipient samples were collected for sequencing: (*i*) 200 µl of larval water from each replicate flask, collected when larvae had molted to the third instar, (*ii*) third instar larvae (pools of 5) collected from the same flasks, and at least three individual sugar-fed adult females that had emerged from pupae either (*iii*) three or (*iv*) six days prior to collection. Pupae produced from replicate flasks containing the same donor pool were pooled and transferred to sterile water in a sterile plastic chamber for adult emergence. Newly emerged adults were provided 10% sucrose in sterile water *ad libitum* until collection as described above. Total genomic DNA was extracted from all recipient samples using a NucleoSpin Tissue Kit (Machery-Nagel, Düren, Germany). Larvae and adults were surface sterilized as described above before DNA isolation.

### Amplicon library construction, sequencing, and data analysis

Subsamples from the DNA extracts were used for amplifying the V3-V4 region of the bacterial 16S rRNA gene using primers 341F (CCTACGGGNGGCWGCAG) and 805R (GACTACHVGGGTATCTAATCC) [59], followed by PCR amplification for Illumina sequencing. Samples were paired-end sequenced (2 × 250-bp) on an Illumina MiSeq. Sequence reads were processed using the DADA2 pipeline in QIIME 2-2019.4 [60, 61]. In brief, sequence reads were first filtered using DADA2’s recommended parameters and an expected error threshold of 0.5. Filtered reads were then de-replicated and denoised using default parameters. After building the ASV table and removing chimeras and low-abundance (<0.005%) ASVs, taxonomy was assigned using a Naive Bayes classifier natively implemented in QIIME 2-2019.4 and pre-trained against the Greengenes reference database (13.8) [62, 63]. A phylogenetic tree was built using FastTree (v2.1.3) [64] from a multiple sequence alignment made with the MAFFT alignment tool [65] against the Greengenes Core reference alignment [66]. Patterns of diversity within the ASV table were analyzed using standard workflows in QIIME 2-2019.4, with a sampling depth of 1,000 reads per sample and default parameters. Downstream statistical analyses were performed using R (http://www.r-project.org/).

## Results

### Adult mosquitoes harbor relatively complex bacterial communities that can be isolated for transplantation

For each donor species (*Ae. aegypti* and *Cx. quinquefasciatus*), we characterized the microbiota of 40 individual adult mosquitoes by sequencing the V3-V4 regions of the 16S rRNA gene. After filtering, denoising, merging, and removing chimeras, we obtained a total of 2,544,303 reads with a median sequencing depth of 32,803 reads per sample (Additional file 1). We obtained an unusually low number of reads (<1,000) for a single *Cx. quinquefasciatus* sample (Additional file 1), which was removed from subsequent analyses. We then assigned taxa and plotted relative abundance of taxa across samples.

We identified 103 and 120 ASVs across all of the *Ae. aegypti* and *Cx. quinquefasciatus* individuals we sampled, respectively (Additional file 2). However, the vast majority of our sequencing reads (>90%) were assigned to ASVs belonging to one of four genera: *Serratia, Asaia, Cedecea* and another, unclassified genus within the *Enterobacteriaceae* (Fig. 2a; Additional file 2). Considering both the presence/absence and relative abundance of all of the ASVs we detected, bacterial communities present in both donor species exhibited significant differences in both alpha diversity (as measured by Shannon’s H index and ASV richness) (Fig. 2b; Additional file 1) and beta diversity (as measured by the Bray-Curtis dissimilarity index) (Fig. 2c), which were associated with shifts in the relative abundance of specific community members detected in each species (Fig. 2a; Additional file 2). As expected, these included the presence of *Wolbachia* in *Cx. quinquefasciatus* donor individuals and the near complete absence of *Wolbachia* across the *Ae. aegypti* donor individuals we sampled, which is consistent with no established natural infection ever being observed in this species [67, 68]. *Cx. quinquefasciatu*s donor individuals further contained a notably greater percentage of taxa within the genus *Serratia*, while *Ae. aegypti* donor individuals contained a greater percentage of *Asaia* spp. (Fig. 2a; Additional file 2). Interestingly, we also detected significant negative correlations between the relative abundance of ASVs belonging to the genus *Serratia* and ASVs belonging to other taxa within the family *Enterobacteriaceae* across all *Ae. aegypti* and *Cx. quinquefasciatus* donors we sampled (Additional files 2 & 3), which is consistent with previous results in field-collected mosquitoes of the same species and our recent experimental findings [52, 57].

**Figure 2.**
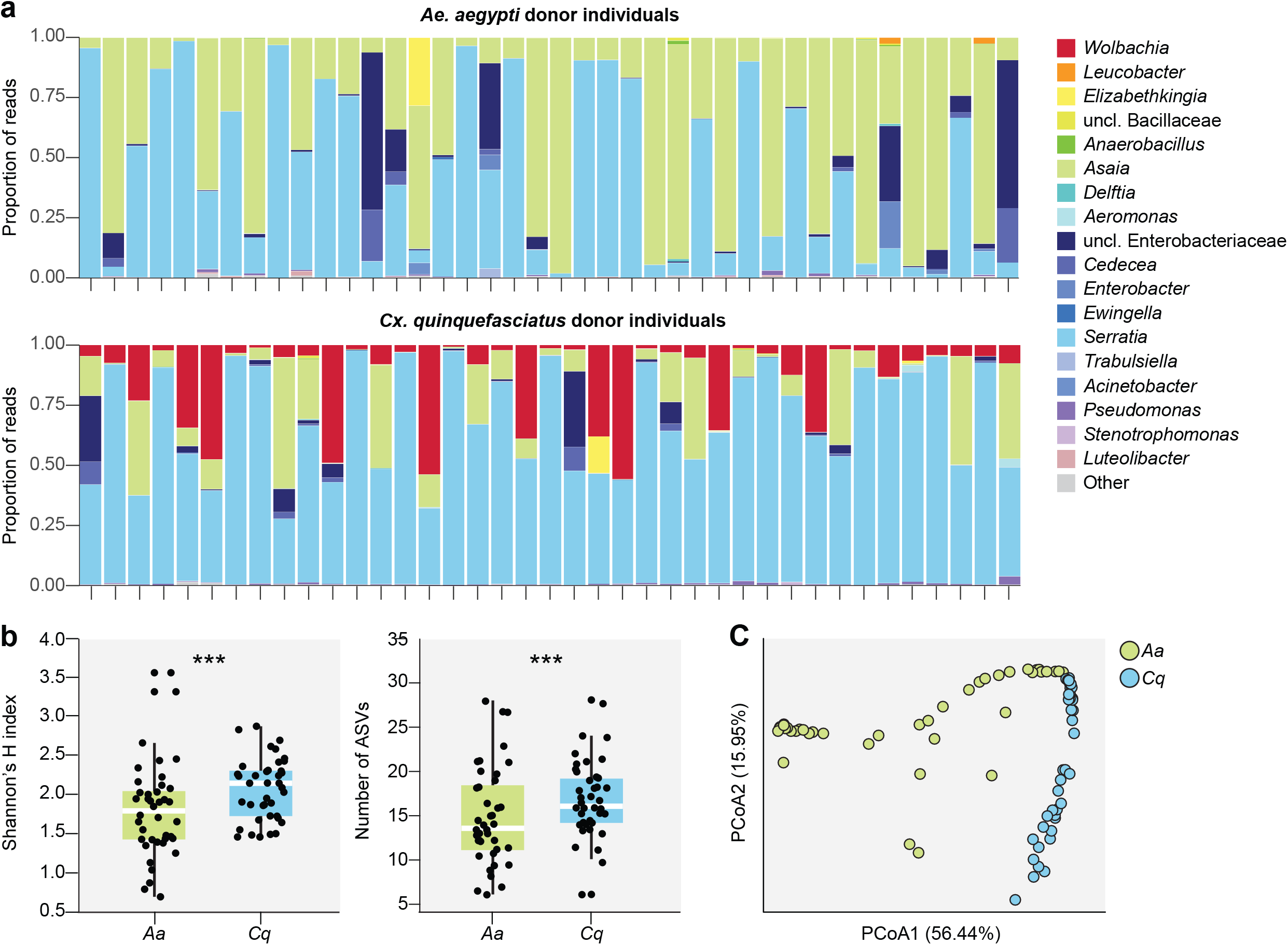
**a** Relative abundance of bacterial genera in individual adult mosquitoes sampled from conventionally maintained populations of each of the donor species used in the study (*i*.*e*., *Ae. aegypti* and *Cx. quinquefasciatus*). Adults were provided 10% sucrose in water (wt/vol) *ad libitum* prior to being sampled 3-4 days post-emergence. Each bar represents an individual mosquito. Low abundance genera (<1%) are represented by the ‘Other’ category. **b & c** Alpha and beta diversity of *Ae. aegypti* (*Aa*) and *Cx. quinquefasciatus* (*Cq*) donor individuals. Panel **b** shows the difference in alpha diversity between *Ae. aegypti* and *Cx. quinquefasciatus* donor individuals as measured by Shannon’s H index (left) and ASV richness (right). Box-and-whisker plots show high, low, and median values, with lower and upper edges of each box denoting first and third quartiles, respectively. Asterisks (***) indicate significant differences between *Ae. aegypti* and *Cx. quinquefasciatus* donor individuals (Kruskal-Wallis test, P < 0.05). Panel **c** shows the difference in beta diversity between *Ae. aegypti* and *Cx. quinquefasciatus* donor individuals. Principal coordinates analysis using the Bray-Curtis dissimilarity index identified significant clustering by donor species (PERMANOVA, P = 0.001).

In order to determine how many individuals were required to transfer a representative microbiome to recipients, we generated homogenates using pools of 10, 20, 40, or 80 *Ae. aegypti* or *Cx. quinquefasciatus* donor individuals and assessed the diversity of ASVs recovered in each pool relative to ASV diversity across the entire donor species populations using 16S rRNA gene amplicon sequencing. We obtained a total of 438,839 sequences representing 68 and 64 ASVs across the four *Ae. aegypti* and four *Cx. quinquefasciatus* donor pools we sequenced, respectively, and each pool was dominated by the same four genera detected in the subset of individuals we sequenced from each of our donor species populations (Fig. 3a; Additional files 1 & 2). Each pool also captured the majority of donor ASV diversity (>96%), although recovery varied with respect to how common a given ASV was across all of the donor individuals we sequenced (Fig. 3b; Additional file 2). The number of individuals used to generate each pool had the greatest impact on recovery of rare ASVs (*i*.*e*., those present in <50% of the donor individuals we sequenced), with significantly fewer rare ASVs being recovered in the pool generated using 10 individuals. However, there were no significant differences in recovery of more common ASVs, which were present in ε50% of donor individuals and constituted >93% of individual donor sequences, between any of the pools we generated (Fig. 3b; Additional file 2).

**Figure 3.**
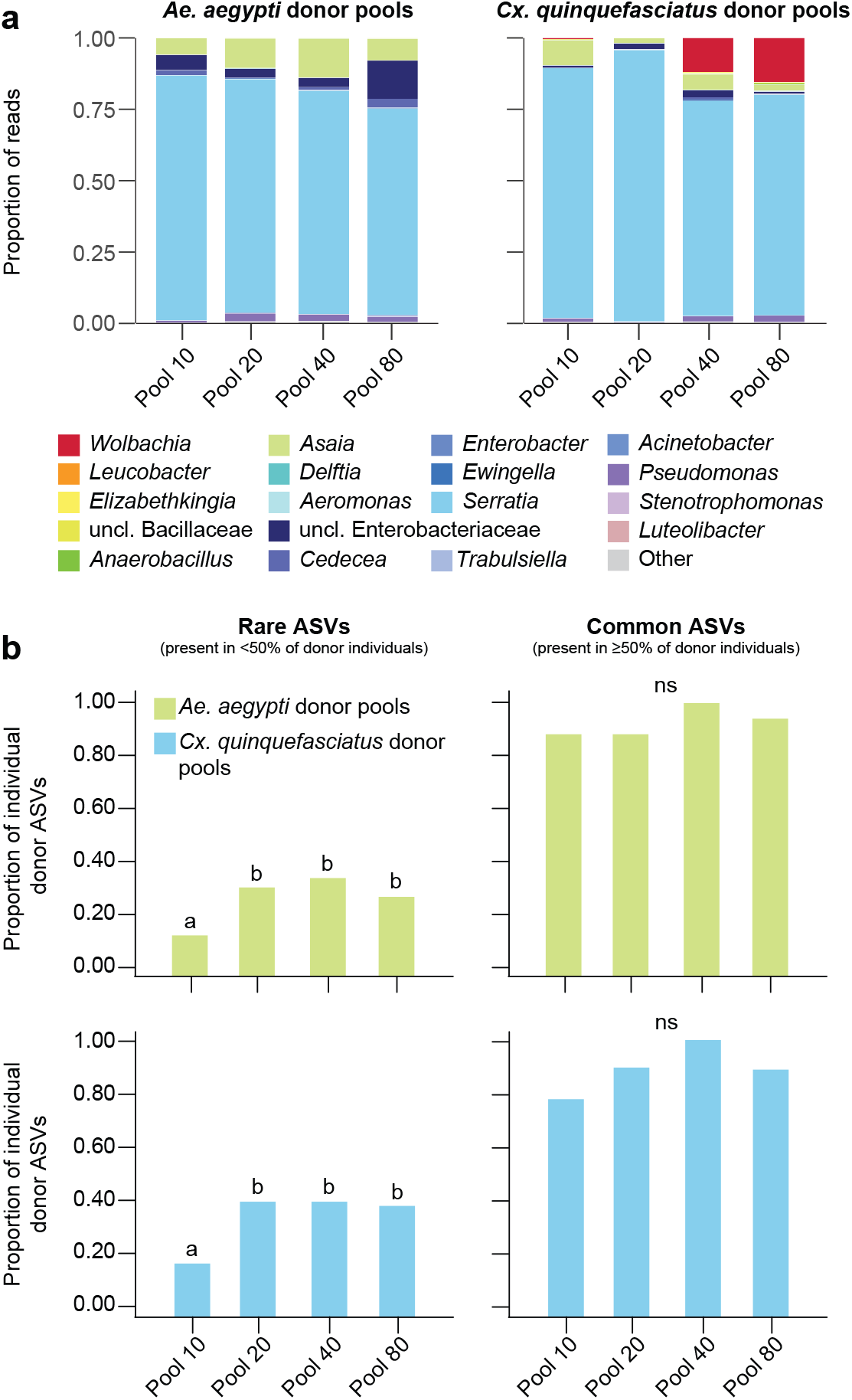
**a** Relative abundance of bacterial genera in *Ae. aegypti* and *Cx. quinquefasciatus* donor pools. Low abundance genera (<1%) are represented by the ‘Other’ category. **b** Proportion of rare ASVs (left) and common ASVs (right) found in at least one donor individual that were present in a given donor pool. An ASV was considered “rare” if it was detected in <50% of donor individuals, while an ASV was considered “common” if it was detected in ≥50% of donor individuals. Pools that do not share a letter above the bars had significantly different results as determined by paired Fisher’s Exact tests (P < 0.05); ns, pools not significantly different (P > 0.05).

### Microbiota transplantation recapitulates donor microbial diversity in recipient individuals

To assess our ability to transplant microbiota between different mosquito species, we introduced each of our *Ae. aegypti* and *Cx. quinquefasciatus* donor pools into the water of cultures containing axenic *Ae. aegypti* larvae. These cultures were subsequently maintained under standard rearing conditions, and 16S rRNA gene amplicon sequencing was used to assess bacterial diversity in replicate samples of the aquatic habitat (water in rearing flasks), larvae (collected as third instars), and adults from each culture (collected 3- and 6-days post-emergence). A total of 50, 40, and 108 water, larval, and adult samples were sequenced, respectively, with a total of five samples being discarded prior to downstream analyses due to low sequencing depth (<1,000 reads) (Additional file 1).

The resulting dataset, which contained a total of 4,540,617 sequences across all of the recipient water, larval, and adult samples we sequenced (Additional file 1), revealed that 64 and 61 of the 68 and 64 ASVs found in *Ae. aegypti* and *Cx. quinquefasciatus* donor pool communities, respectively, representing >99% of all donor pool sequences, were detected in recipient samples (Additional file 2). The majority of these ASVs (>70%) also persisted in recipient communities over time, although there were dramatic shifts in both alpha and beta diversity across the different life stages and water samples we sequenced (Fig. 4 & Fig. 5; Additional files 1, 2; 4 & 5). Alpha diversity (as measured by Shannon’s H index and ASV richness) was overall highest in water and larvae, while adult recipient individuals harbored communities that did not significantly differ in alpha diversity from input donor communities (Fig. 4b; Additional file 1), regardless of the size of the input donor pool (Additional files 1, 4 & 5). Differences in beta diversity, measured as average Bray-Curtis dissimilarity, were also overall higher between input donor communities and the recipient water and larval samples we collected than between input donor communities and recipient adult samples (Fig. 5), although the degree of similarity between recipient adult and input donor communities varied over time post emergence and as a function of the size of the input donor pool (Additional file 6). More specifically, recipient water and larval communities displayed striking amplification of taxa within the Chitinophagaceae, Microbacteriaceae (*Leucobacter* and *Microbacterium* spp.), and Flavobacteriaceae (*Flavobacterium, Chryseobacterium*, and *Elizabethkingia* spp.) that were less abundant in input donor communities (Fig. 4a; Additional file 2). In contrast, adult recipient communities displayed striking amplification of taxa within the genera *Serratia, Asaia, Cedecea*, and an unclassified genus within the *Enterobacteriaceae*, consistent with input donor communities (Fig. 4a; Additional file 2).

**Figure 4.**
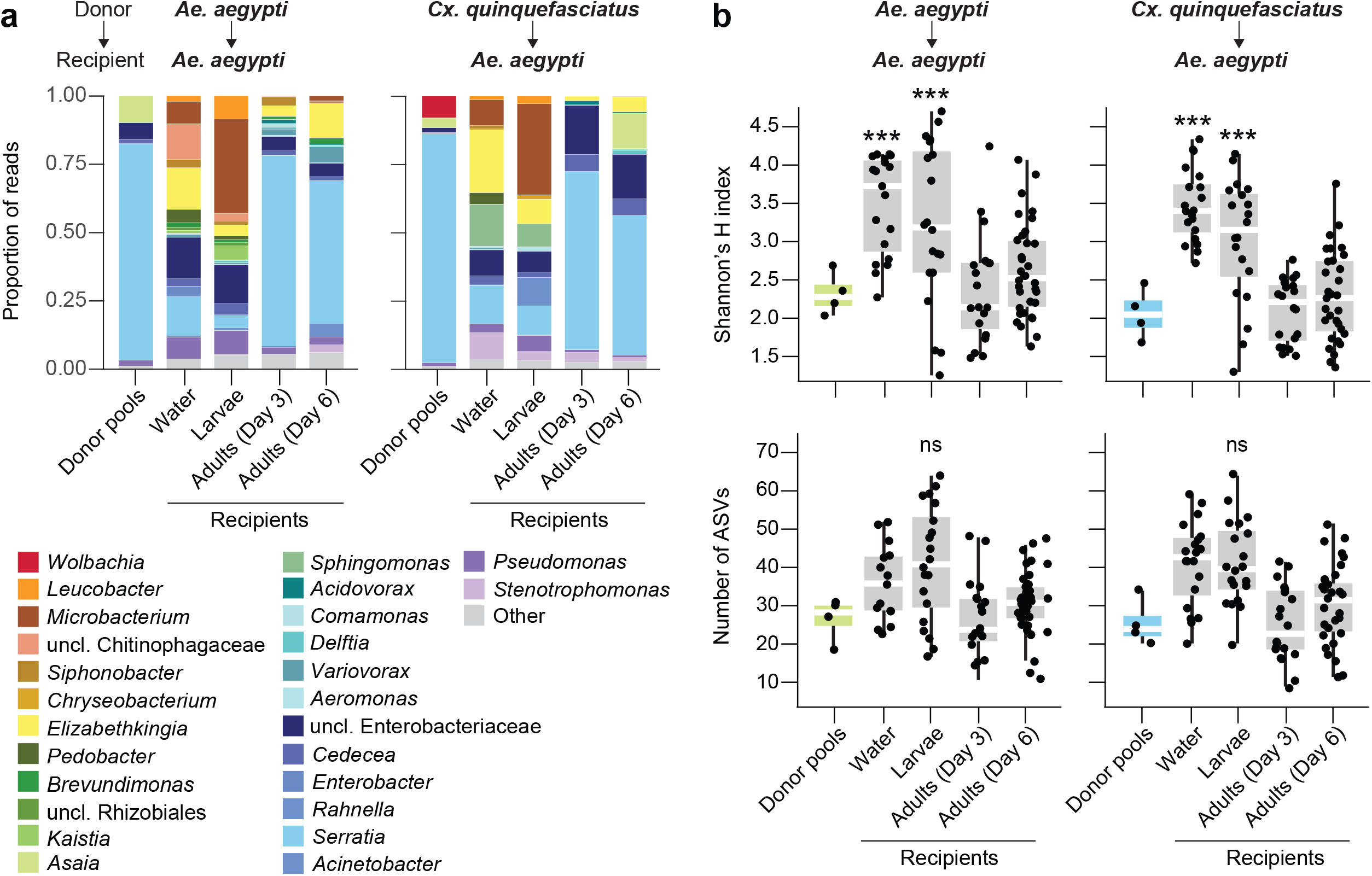
**a** Relative abundance of bacterial genera in donor pools and recipient samples. Biological replicates were pooled for the bar graphs presented. Low abundance genera (<1%) are represented by the ‘Other’ category. **b** Alpha diversity of donor pools and recipient samples, as measured by Shannon’s H index and ASV richness. Box-and-whisker plots show high, low, and median values, with lower and upper edges of each box denoted first and third quartiles, respectively. Asterisks (***) indicate samples that significantly differed from the donor pools (Dunn’s test with Bonferroni correction, P < 0.0125).

**Figure 5.**
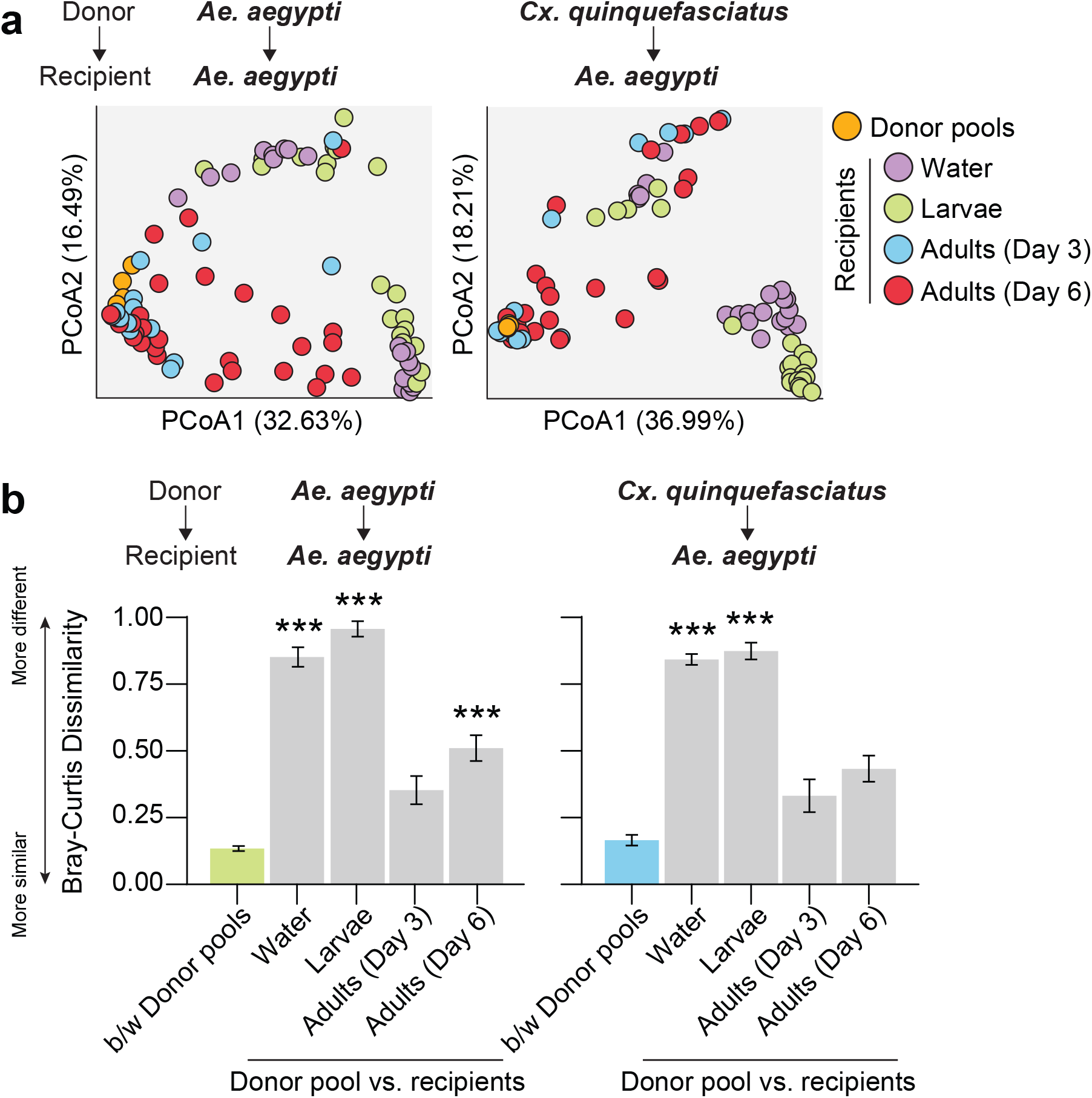
**a** Principal coordinates analysis using the Bray-Curtis dissimilarity index. Circles are colored by sample source (donor pools: orange, water: purple, larvae: green, 3-day old adults: blue, 6-day old adults: red). **b** Average Bray-Curtis dissimilarity between (b/w) donor pools versus between a given donor pool and recipient samples. Mean values ± standard errors are shown. Asterisks (***) indicate comparisons for which the average dissimilarity between a given donor pool and group of recipient samples was significantly higher than that expected as a result of the transplantation procedure itself (*i*.*e*., between donor pools) (Dunn’s test with Bonferroni correction, P < 0.0125).

### Both host and environmental factors impact microbiota transplantation efficacy

To assess whether microbiota transplantation recapitulated observed differences in microbiota diversity between donor species in recipient individuals, we last compared alpha and beta diversity of donor *Ae. aegypti* and *Cx. quinquefasciatus* individuals to recipient *Ae. aegypti* adults emerging from cultures inoculated with pools generated from each donor species. Consistent with our previous results comparing bacterial communities in donor individuals of each species (Fig. 2), communities in recipient adults that emerged from cultures inoculated with pools generated from each donor species exhibited significant differences in both alpha diversity (as measured by Shannon’s H index and ASV richness) (Fig. 6b) and beta diversity (as measured by the Bray-Curtis dissimilarity index) (Fig. 6c). However, these differences were associated with shifts in the relative abundance of community members that were relatively rare in input donor communities (*e*.*g*., taxa within the genera *Elizabethkingia, Acinetobacter, Pseudomonas*, and *Stenotophomonas*) (Fig. 2 & Fig. 6a; Additional file 2). More specifically, the predominance of *Asaia* in *Ae. aegypti* donor communities was not observed in recipient individuals, likely owing to the inability of this bacterium to reliably persist in the water of larval cultures (Fig. 6a & Additional file 2). *Wolbachia* from *Cx. quinquefasciatus* donor communities did not infect recipient individuals (Fig. 6a & Additional file 2), although this was expected given that transfections of *Wolbachia* into mosquitoes requires access through the germline. While these patterns were generally similar between groups of recipient individuals inoculated with input donor pools generated using different quantities of mosquitoes, adult recipients inoculated with pools generated using fewer individuals exhibited more variable bacterial communities overall, due to the stochastic proliferation of rare donor taxa and/or antagonism of *Serratia* by *Enterobacteriaceae*, consistent with our previous observations across donor individuals (Fig. 6a; Additional files 2 & 7).

**Figure 6.**
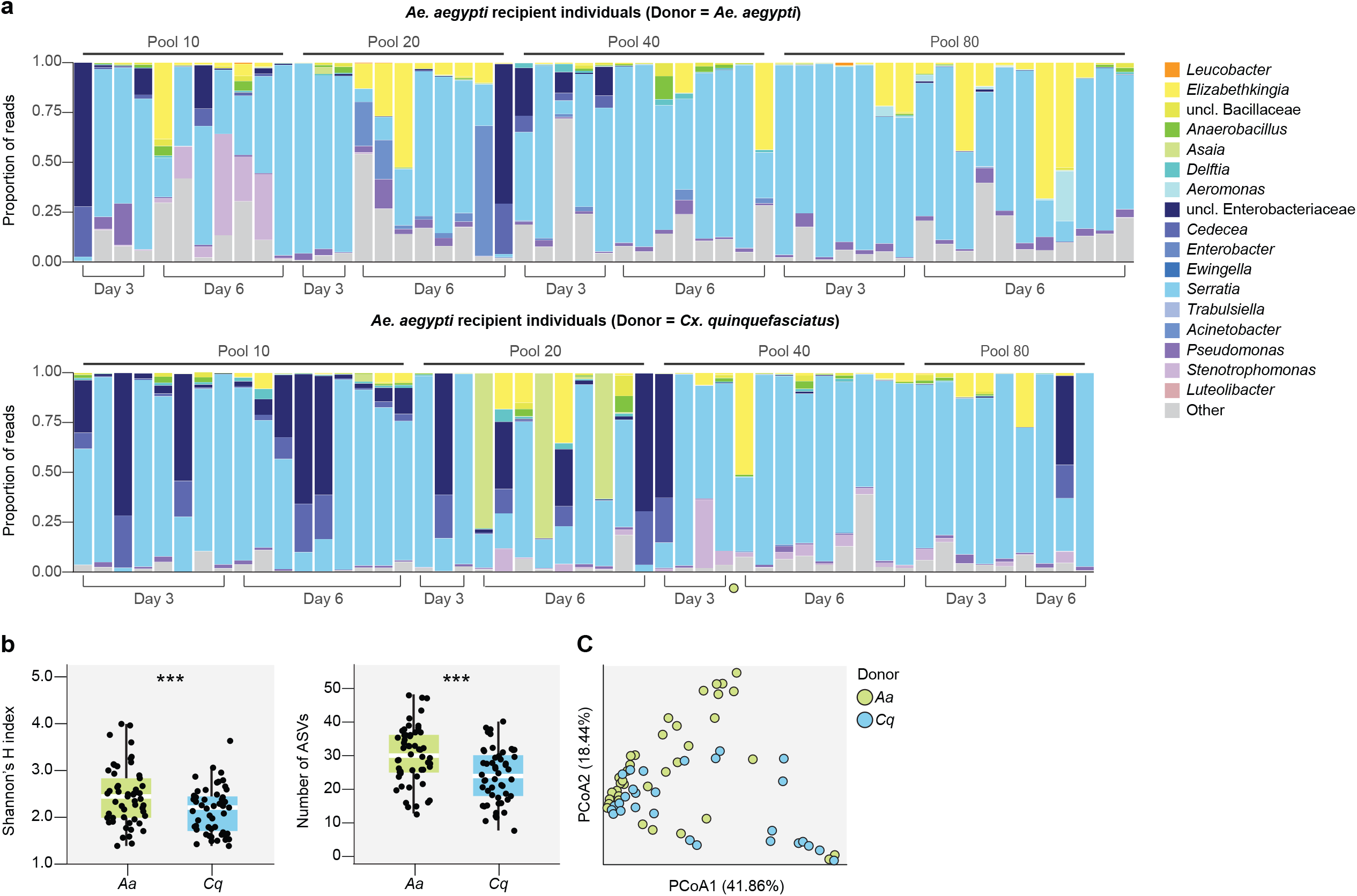
**a** Relative abundance of bacterial genera in 3-day and 6-day old *Ae. aegypti* recipient adults that emerged from cultures inoculated with donor pools generated from either *Ae. aegypti* or *Cx. quinquefasciatus* individuals. Each bar represents an individual mosquito. Genera representing >1% of reads from any one donor individual are listed in the legend; all other taxa are grouped together under ‘Other’. **b & c** Alpha and beta diversity of recipient *Ae. aegypti* adults emerging from cultures inoculated with pools generated from *Ae. aegypti* (*Aa*) or *Cx. quinquefasciatus* (*Cq*) donor individuals. Panel **b** shows the difference in alpha diversity between *Aa* and *Cq* recipient adults as measured by Shannon’s H index (left) and ASV richness (right). Box-and-whisker plots show high, low, and median values, with lower and upper edges of each box denoting first and third quartiles, respectively. Asterisks (***) indicate significant differences between *Aa* and *Cq* recipient adults (Kruskal-Wallis test, P < 0.05). Panel **c** shows the difference in beta diversity between *Aa* and *Cq* recipient adults. Principal coordinates analysis using the Bray-Curtis dissimilarity index identified significant clustering by donor species (PERMANOVA, P = 0.001).

## Discussion

Numerous studies describing the diversity and function of mosquito-associated microbial taxa and assemblages have been published recently, leading to the discovery of potential interactions between mosquitoes and their microbiota that could be manipulated to enhance or reduce mosquito fitness and/or vector competency (reviewed in [36]). However, questions remain about the reproducibility, applicability, and physiological relevance of these data owing to discrepancies in experimental technique, lack of standardization, and the use of laboratory-colonized mosquito strains and species that harbor microbiota that vary substantially within and between different laboratories and that differ substantially from naturally occurring mosquitoes in the field [12, 13, 69-74]. In this study, we developed a simple approach to successfully isolate donor microbial communities from adult mosquitoes. We then used this approach to transfer microbiota within (*i*.*e*., *Ae. aegypti* donors to *Ae. aegypti* recipients) and between (*i*.*e*., *Cx. quinquefasciatus* donors to *Ae. aegypti* recipients) different donor and recipient species in order to assess our ability to recapitulate donor microbial diversity in adult recipients. We selected these donor and recipient species for two reasons. First, both *Ae. aegypti* and *Cx. quinquefasciatus* are biomedically relevant vector species that are commonly studied in the laboratory [7]. Previous studies have also carefully characterized microbiota acquisition and assembly across *Ae. aegypti* life history [10]. Second, while *Cx. quinquefasciatus* mosquitoes are readily infected by *Wolbachia* in the laboratory and commonly exhibit natural infections in the field [75 76], *Ae. aegypti* mosquitoes are generally recalcitrant to infection and are not known to harbor natural infections [67, 68]. By introducing microbiota derived from both *Ae. aegypti* and *Cx. quinquefasciatus* donors into *Ae. aegypti* recipients, we could therefore also establish whether our transplantation approach (*i*) recapitulated patterns of microbiota assembly and persistence previously observed in *Ae. aegypti* (positive control), and (*ii*) supported the expectation that we would not be able to successfully transfer *Wolbachia* from *Cx. quinquefasciatus* donors to *Ae. aegypti* recipients (negative control).

Our results demonstrate the successful transfer of bacteria from donor to recipient populations of mosquitoes, with recipient adult individuals retaining the vast majority of donor bacterial diversity. Our results also demonstrate that relatively small pools of donor microbiota (*i*.*e*., derived from ≥10 donor individuals) are sufficient to capture taxa that are highly abundant and/or common in a given donor mosquito population. This suggests that our methods could readily be applied to study microbiota isolated from field populations, even in cases where large-scale mosquito collections are difficult. However, future work will be necessary to confirm that the patterns observed using our approach are indeed the same for donor microbiota generated from field mosquitoes that harbor microbial communities that are more diverse than those present in laboratory mosquitoes and that are not adapted to the laboratory environment [12, 13, 69-74]. Recently developed methods to generate axenic adults [28, 53-55] also strongly position us to examine the potential to adapt our protocol to introduce field-derived donor microbiota pools directly into adults via a sugar meal and therefore avoiding selective pressures exerted on introduced microbes by standard laboratory larval diets and larval growth and molting.

These results also highlight the value of our approach for studying the underlying bases of mosquito-microbe interactions, as has been demonstrated in other animals. For example, microbiome transplantation approaches have been employed in *Nasonia* wasps to assess the selective pressures contributing to observed patterns of phylosymbiosis in the system [41] as well as *Drosophila melanogaster* flies and *Bombus terrestris* bumblebees to identify relationships between host microbiota composition and different thermotolerance or immunity phenotypes [39, 40]. Reciprocal microbiome transplantations have also been performed across different vertebrate animal species. For example, reciprocal microbiota transplantations between zebrafish and mice have revealed factors specific to the host gut habitat that shape microbial community structure in each species [38]. Indeed, our results point to both host and environmental factors in shaping microbiota acquisition and persistence in mosquitoes. That the microbiota in recipient adult individuals looked most similar to the microbiota of donor individuals supports previous studies in *Ae. Aegypti* identifying life stage as a dominant factor shaping mosquito microbiota [10, 11, 14, 69, 70, 77-82]. Additionally, there were differences in microbiota diversity between our donor species in recipient adults which supports previous findings indicating that while community membership may be largely driven by the environment and life stage, community features such as total and taxon-specific abundances may also be shaped by host genetics [83, 84]. Future studies could employ microbiome transplantation to improve our understanding of the factors shaping microbiome acquisition and assembly in mosquitoes and the mechanisms by which specific microbial taxa and assemblages contribute to different mosquito traits under controlled conditions.

Future work is also warranted to determine if the approach developed herein is relevant for other microbes such as fungi or viruses, which are also known to impact mosquito biology [85, 86]. The applicability of this approach for other microbes likely depends on their biology, and in particular, their mode of transmission. For example, extracellular members of the mosquito gut microbiota that are commonly detected in environments where mosquitoes persist in the laboratory are likely to be transferred, while those that are intracellular may not be. In line with this, our results supported the expectation that the intracellular bacterium *Wolbachia*, which was present in the *Cx. quinquefasciatus* donors used in this study, would not be successfully transferred to *Ae. aegypti* recipients. We also appreciate that tissue localization of bacteria and other microbes may be pertinent for the transplantation process. Here, we prepared donor microbiota from whole-body mosquitoes and similarly assessed microbiota assembly and persistence in whole-body recipient individuals. Additional work will be necessary to establish patterns of donor microbiota colonization across recipient host tissues and/or to validate methods for isolation of microbiota from specific donor tissues. Methods for long-term preservation of donor microbiota will also be necessary to facilitate long-term studies and intra- and inter-laboratory comparisons of microbiota assembly across different host strains. Nevertheless, our results provide a critical first step toward the standardization of microbiome studies in the field of vector biology to include mosquito hosts colonized by defined microbiota that can be replicated within and between labs. They also provide a critical first step toward our ability to recapitulate and study field-derived microbiota in laboratory settings.

## Conclusions

We have successfully isolated and transplanted microbiomes from donor to recipient mosquitoes. This approach lays the foundation for future work to facilitate standardized studies of mosquito-microbe interactions, examine host-microbe interactions, and to identify effective strategies for manipulating mosquito microbiota to control mosquito populations and mosquito-borne diseases.

## List of Abbreviations

Not applicable.

## Declarations

### Ethics approval and consent to participate

Not applicable.

### Consent for publication

Not applicable.

### Availability of data and materials

Raw Illumina reads are available in the NCBI Sequence Read Archive (http://www.ncbi.nlm.nih.gov/sra) under BioProject ID PRJNA767109.

### Authors’ contributions

KLC, SH, and GLH conceived and designed the experiments. SH performed the experiments. KLC carried out the data analysis. KLC wrote the initial manuscript, and SH and GLH contributed to revisions.

## Acknowledgements

We thank George Golovko, Kamil Khanipov, and Maria Pimenova of the Department of Pharmacology and Toxicology, Sealy Center for Structural Biology, University of Texas Medical Branch (Galveston, TX USA) for assistance with sequencing. We also thank the insectary staff at the University of Texas Medical Branch for assistance in rearing mosquitoes.

## Additional files

**Additional file 1: Supplementary Table 1**. Sequencing and diversity statistics for 16S rRNA gene amplicon libraries prepared from donor and recipient samples.

**Additional file 2: Supplementary Table 2**. Taxonomic assignments and prevalence of each ASV in each sample, along with whether a particular ASV was considered ‘rare’ or ‘common’ among individuals of a particular donor species.

**Additional file 3: Supplementary Figure 1**. Significantly negative correlation between relative abundance of ASVs belonging to the bacterial family Enterobacteriaceae and ASVs belonging to the genus *Serratia* across all *Ae. aegypti* and *Cx. quinquefasciatus* donor individuals (Spearman’s rank test, P < 0.05).

**Additional file 4: Supplementary Figure 2**. (Left) Relative abundance of bacterial genera in *Ae. aegypti* donor pools and recipient samples. Biological replicates were pooled for the bar graphs presented. Low abundance genera (<1%) are represented by the ‘Other’ category. (Right) Alpha diversity of donor pools and recipient samples, as measured by Shannon’s H index and ASV richness. Box-and-whisker plots show high, low, and median values, with lower and upper edges of each box denoting first and third quartiles, respectively. Asterisks (***) indicate samples that significantly differed from the donor pools (Dunn’s test with Bonferroni correction, P < 0.0125).

**Additional file 5: Supplementary Figure 3**. (Left) Relative abundance of bacterial genera in *Cx. quinquefasciatus* donor pools and recipient samples. Biological replicates were pooled for the bar graphs presented. Low abundance genera (<1%) are represented by the ‘Other’ category. (Right) Alpha diversity of donor pools and recipient samples, as measured by Shannon’s H index and ASV richness. Box-and-whisker plots show high, low, and median values, with lower and upper edges of each box denoting first and third quartiles, respectively. Asterisks (***) indicate samples that significantly differed from the donor pools (Dunn’s test with Bonferroni correction, P < 0.0125).

**Additional file 6: Supplementary Figure 4**. Average Bray-Curtis dissimilarity between (b/w) donor pools versus between a given donor pool and recipient samples. Mean values ± standard errors are shown. Asterisks (***) indicate comparisons for which the average dissimilarity between a given donor pool and group of recipient samples was significantly higher than that expected as a result of the transplantation procedure itself (*i*.*e*., between donor pools) (Dunn’s test with Bonferroni correction, P < 0.0125).

**Additional file 7: Supplementary Figure 5**. Significantly negative correlation between relative abundance of ASVs belonging to the bacterial family Enterobacteriaceae and ASVs belonging to the genus *Serratia* across all recipient *Ae. aegypti* adults (Spearman’s rank test, P < 0.05).

