## Supplementary figures and images for "Inter-species microbiota transplantation recapitulates microbial acquisition and persistence in mosquitoes"

### Supplemental Figure 1

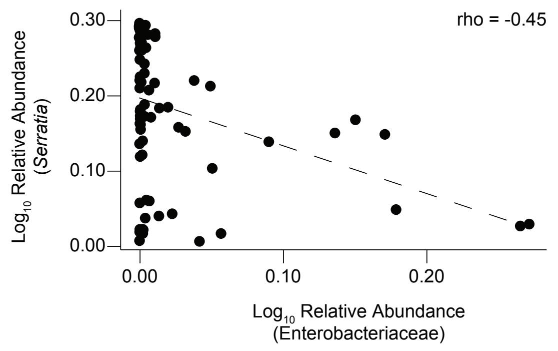

### Supplemental Figure 2

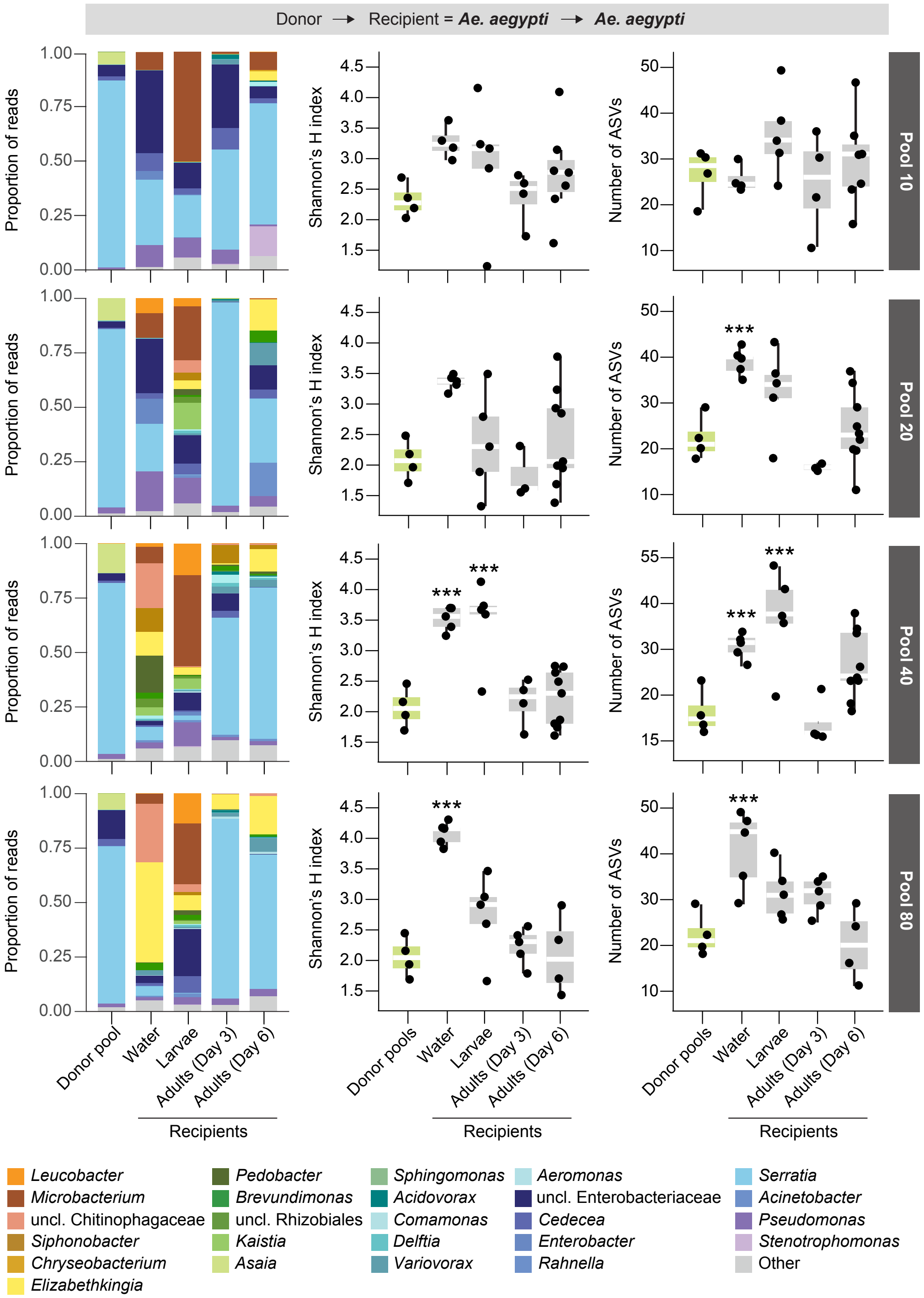

### Supplemental Figure 3

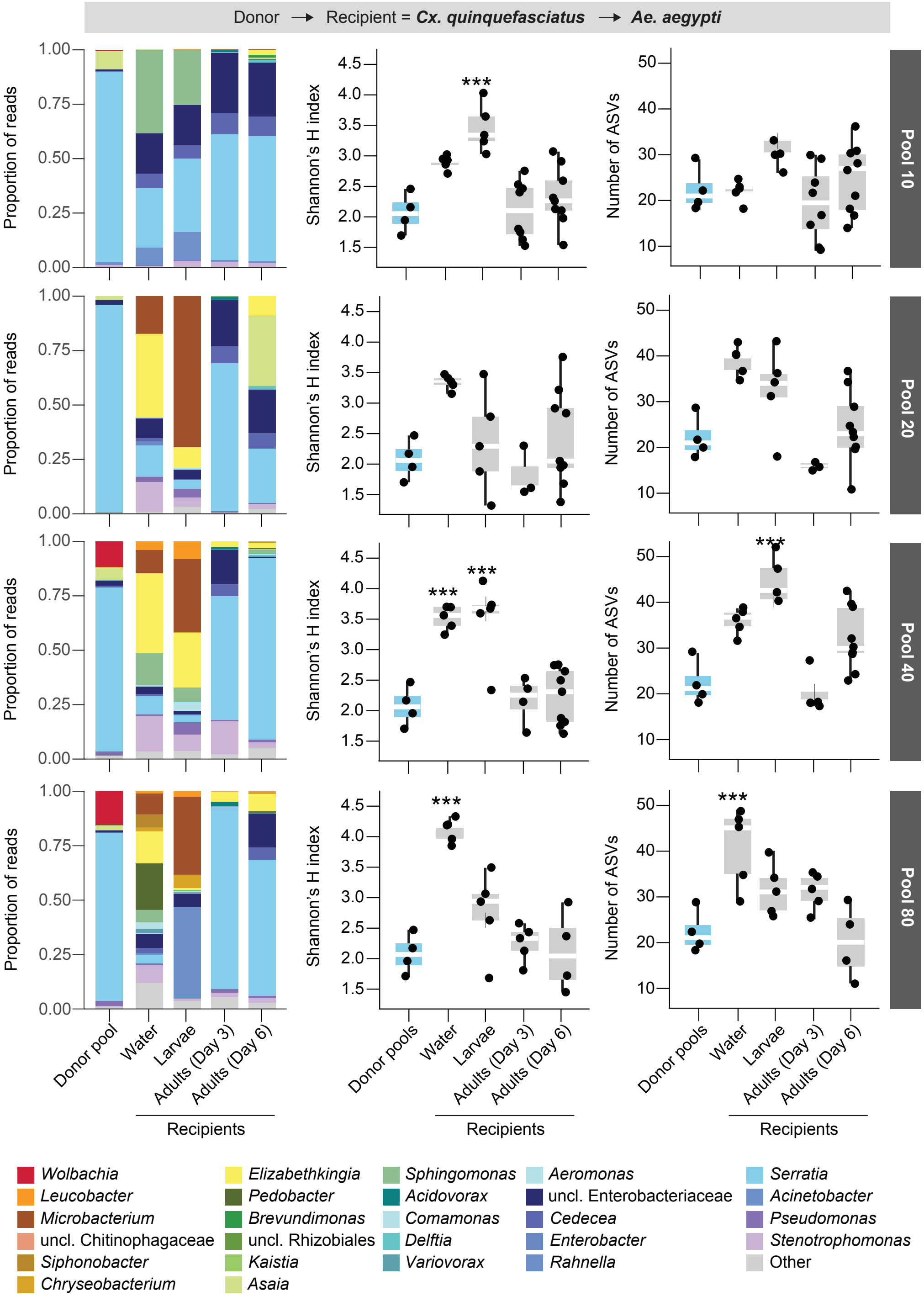

### Supplemental Figure 4

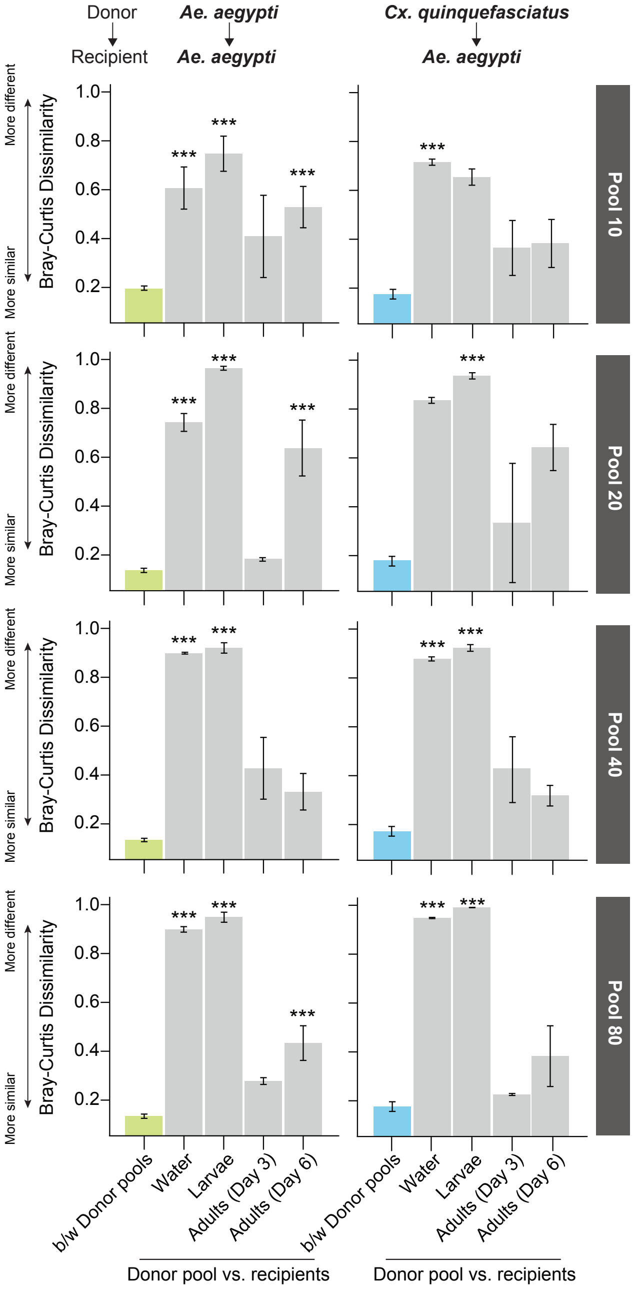

### Supplemental Figure 5

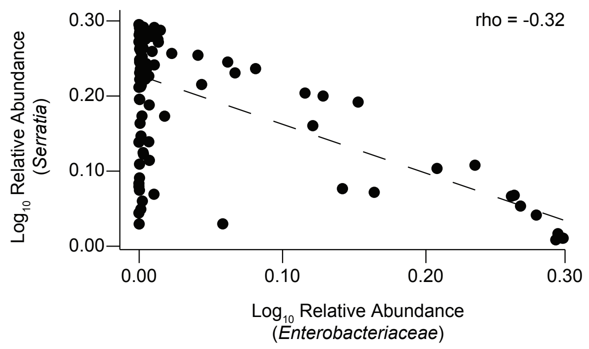
